# Spatio-temporal Memory for Navigation in a Mushroom Body Model

**DOI:** 10.1101/2020.10.27.356535

**Authors:** Le Zhu, Michael Mangan, Barbara Webb

## Abstract

Insects, despite relatively small brains, can perform complex navigation tasks such as memorising a visual route. The exact format of visual memory encoded by neural systems during route learning and following is still unclear. Here we propose that interconnections between Kenyon cells in the Mushroom Body (MB) could encode spatio-temporal memory of visual motion experienced when moving along a route. In our implementation, visual motion is sensed using an event-based camera mounted on a robot, and learned by a biologically constrained spiking neural network model, based on simplified MB architecture and using modified leaky integrate-and-fire neurons. In contrast to previous image-matching models where all memories are stored in parallel, the continuous visual flow is inherently sequential. Our results show that the model can distinguish learned from unlearned route segments, with some tolerance to internal and external noise, including small displacements. The neural response can also explain observed behaviour taken to support sequential memory in ant experiments. However, obtaining comparable robustness to insect navigation might require the addition of biomimetic pre-processing of the input stream, and determination of the appropriate motor strategy to exploit the memory output.

## 1 Introduction

In the task of finding food and going back home, visual memory is one of the key cues used by insects. Ants can use their previous visual experience of a route to follow it efficiently, even when other navigation cues such as path integration are unavailable. Moreover, ants can quickly recognise familiar surroundings after being blown off-course by a gust of wind or displaced by an experimenter [23]. This capability, equivalent to solving the ‘kidnapped robot’ problem, is a recognised challenge for autonomous navigation systems, especially if only monocular visual information is available. Recognising a location from the current visual input can be difficult due to high variability across viewpoints, weather conditions, time of day, and seasons [13], yet insects provide an existence proof that it is possible using low-resolution vision and relatively limited neural processing.

Improved visual recognition has been achieved on robots by matching sequences of images [15, 14] along a route. This reduces the impact of individual mismatches and demonstrates the benefit of incorporating the temporality of visual data in the learning regime. To date, bioinspired navigation models (with the possible exception of [5]) have largely overlooked the potential benefits of temporal cues for visual recognition, instead relying on repeated static view matching (see figure 1), with improvements sought through visual processing schema that remove input variability [20, 22]. Yet it has been observed that the navigation behaviours of ants are not only affected by the current view, but also recent visual experiences [7]. We have recently found that altering the sequence of views experienced by ants on their homeward journey affects recall ability indicating a role for temporal cues (Schwarz et al, under review). Moreover, the peripheral processing in insect vision is highly sensitive to motion, i.e. they fundamentally experience a spatio-temporal input rather than static frames. There is an extensive literature on the use of motion cues by insects [19] for tasks from obstacle avoidance [3], to odometry [4], and identification of camouflaged landmarks [12] during navigation.

**Fig. 1.**
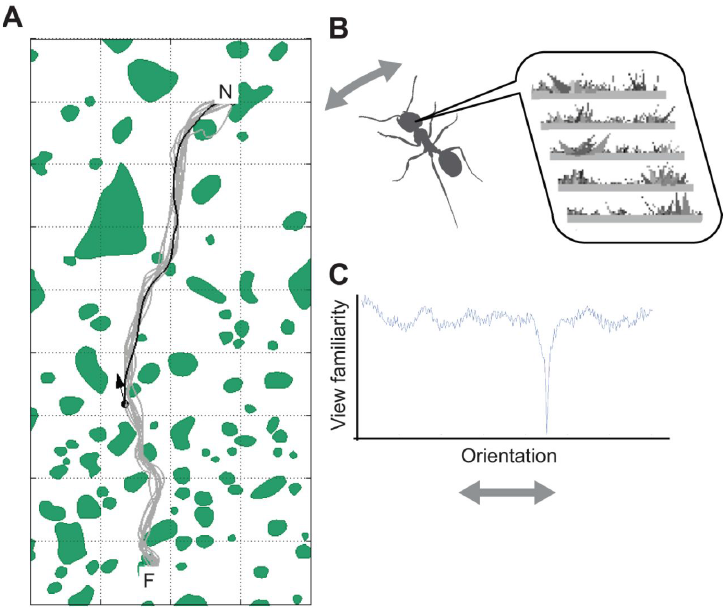
Route following using an image-matching model, e.g. [2]. (A) The ant is assumed to store a set of view snapshots as it moves from the feeder (F) to the nest (N). (B) To follow the route, the ant scans left and right, comparing the current view to all stored views. (C) The view is most familiar (low value) when facing along on the route. Figure copied from [7], with permission.

In this work, we investigate whether sensing and learning of motion flow patterns in a biologically-constrained associative network are sufficient for storage and recall of visual routes.

## 2 Methods

### 2.1 Input: Bio-inspired event-based camera

The input to our system is visual motion delivered in the form of spikes from an event-based camera DAVIS 346 (346×260 pixels). Event-based cameras (also called dynamic vision sensor (DVS)) output a spike when an individual pixel detects brightness change. While conventional cameras capture from the entire sensor at discrete time steps, event-based cameras only transmit pixel data when the intensity of a given pixel shifts up or down by a predetermined amount. This enables a time resolution of up to 1 ns (see figure 2). Event-based cameras are inspired by that of vertebrate retina [17], but the principle of function is also appropriate to represent motion perception in insect eyes. The asynchronous pixels resemble transient photoreceptor responses encoding intensity change; and neurons encoding either ON (dark to bright) or OFF (bright to dark) changes are found in the insect lamina (neurons L1, L2 and L3) [18]. However, the spatial acuity of insect eyes is usually between 2-5 degrees of their visual field, which is lower than our camera resolution. Therefore, before feeding the camera output to the neural network, the event flow is re-sampled by 16*pixel* × 10*pixel* × 1*ms* spatio-temporal windows, with a threshold for activation that also reduces noise. After re-sampling, the effective spatial acuity is 5 degrees and the temporal resolution is 1ms. Note that there are additional processing steps in the insect visual motion pathway, but we have not included these steps in our modelling so far.

**Fig. 2.**
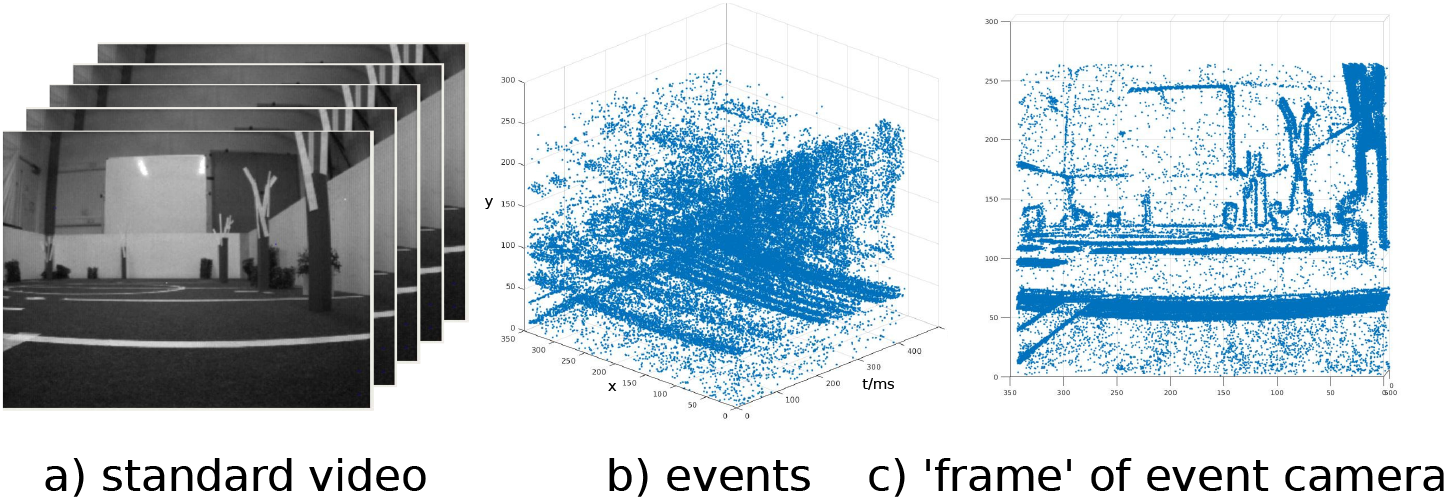
Comparing standard video to an event-based camera view of the test arena in Fig 4. Video (left) has static intensity frames at a fixed rate. ‘Events’ (centre) occur in continuous time whenever a pixel changes intensity; integrated over a period of forward motion, objects are visible in the movement ‘frame’ (right).

### 2.2 Mushroom Body Spiking Neural Network

#### Mushroom Body

The mushroom body (MB) is a multi-sensory processing and learning centre in the insect brain. We have previously implemented an image-matching model based on the MB circuit structure [1]. In that model, visual snapshots were used as input to projection neurons (PN) which fan-out to produce a sparse coding of the pattern by Kenyon Cell (KC) firing activity. Reward signals reduced the connection weights between KCs and an MB output neuron (MBON) for selected patterns corresponding to static images along a route. In the current model, the architecture (figure 3) is similar to [1], but we implement a different learning mechanism, inspired by the unexpected finding [6] that 60% of the input synapses of KCs are from other KCs, and 45% of the KCs output synapses connect to other KCs. Most of the KC-KC connections are axon-axon connections located in the peduncle and MB lobes. We also include the APL neuron which feeds back from the MBON to the KC and provides global inhibition that functions as a gain control. The other key difference is that the PN input is based on the continuous spiking signal from the event-based camera as it is moved along the route.

**Fig. 3.**
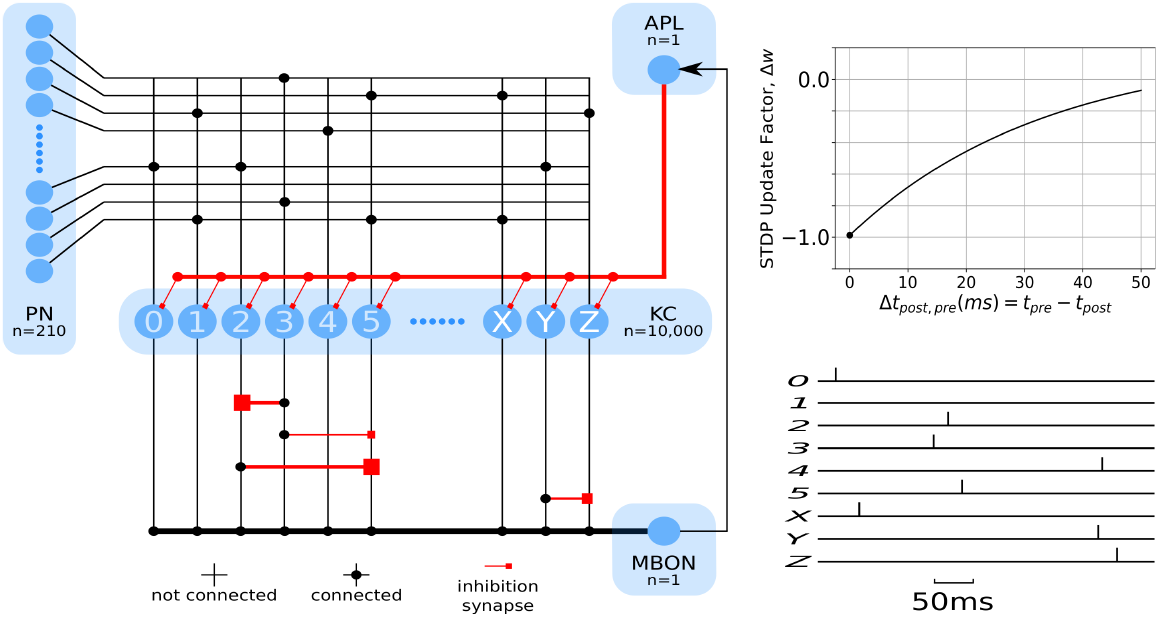
Left: MB network structure (PN: Projection Neuron, APL: Anterior Paired Lateral, MBON: Mushroom Body Output Neuron, KC: Kenyon Cell (from 2 separate lobes, labelled by numbers and letters respectively). Right top: Modified STDP learning rule for KC-KC inhibition. Right bottom: Example of a KC spiking sequence. Here, 3 fires just before 2, followed by 5, so the weights between them change, shown by the varied width of synapses in the network picture. The nearer in time the KC pair fires together, the stronger their inhibitory connection will be.

#### Neuron Model

The neuron models for PN, KC, MBON and APL are all modified from a standard Leaky-Integrated-Fire (LIF) neuron. Modifications are made to match each type of neuron to experimental recorded data as shown in table 2.

The PN is modelled as:

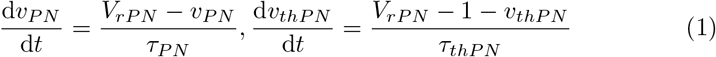

Values for the parameters in this equation can be found in table 1. The two variables *ν*_*PN*_ (PN membrane potential) and *ν*_*thPN*_ (PN threshold voltage) are reset if the membrane potential exceeds threshold *ν*_*thPN*_ :

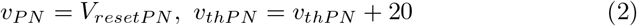

**Table 1.** Neuron model parameters. Values are collected from [11, 21]

| Parameters | PN | KC | MBON | APL |
| --- | --- | --- | --- | --- |
| Resting Potential $V_r$ (mV) | -55 | -60 | -55 | -60 |
| Threshold Voltage $V_{th}$ (mV) | -40 | -45 | 15 | -45 |
| Reset Potential $V_{reset}$ (mV) | -45 | -50 | -50 | -50 |
| Time Constant $\tau$ (ms) | 11.5 | 11.5 | 20 | 11.5 |

**Table 2.** Firing rate (Hz) for MB neuron in recording [9, 16] and our model.

| state \ neuron | PN |  | KC |  | MBON |  |
| --- | --- | --- | --- | --- | --- | --- |
|  | Record | Model | Record | Model | Record | Model |
| Resting | 3.87±2.23 | 4 | 0.025 | 0.03 | N\A | 0 |
| Before Learning | 19.53±10.67 | 20 | 2.32±2.68 | 1.8 | 100–300 | 200–400 |
| After Learning | 19.53±10.67 | 20 | N\A | 1.8 | <100 | <100 |

The adaptive threshold voltage gives each PN a 4Hz baseline firing rate and limits the maximum activity when the input layer is too active. The KC layer has the largest number of neurons. To speed up the network, the model of KC has a constant threshold. Membrane potential of KC (same model for APL) is updated and reset by :

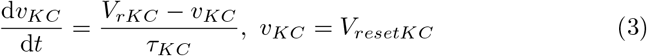

Because the single MBON receives an excitation signal from all 10,000 KCs, the postsynaptic response is changed from membrane potential increase to a synaptic current increase. The membrane potential of MBON is updated by:

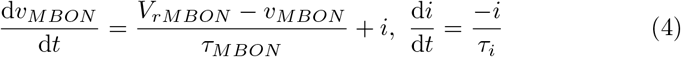

and reset by:

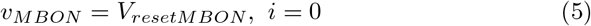

#### Learning Rule

The network architecture and synaptic plasticity learning rule are illustrated in figure 3. The 210 PNs are one-to-one connected to re-sampled camera pixels. Each KC randomly connects to 5-10 PNs. All the parallel axons of KCs terminate at the dendrite of MBON. KCs are divided into two lobes. In each lobe, KCs are fully connected to each other by axon to axon connection. During learning, active KCs will strengthen inhibitory connections to KCs that fire shortly afterwards. Note this is an axon-axon inhibition, i.e., it inhibits the output of the downstream KC onto the MBON, rather than the KC’s activation. The KC-KC synapse weights are all 0 before learning and are altered by STDP. Figure 3 visualises the learned KC-KC synapse based on a firing sequence example. After learning, when the same visual flow occurs, the KC firing sequence will be the same. Excitation from KCs to MBON will be weakened, due to the inhibition generated precisely for this sequence. The output can thus be interpreted as the (un)familiarity between current visual flow and stored memories.

## 3 Experiments and Results

When testing in the indoor environment (shown in figure 4), the event-based camera is carried by a Turtlebot Burger3. The robot was commanded to move forwards at 0.2 m/s, commensurate with the speed of desert ants. In the test arena, visual motion was experienced as the robot followed a straight line from one end to the other.

**Fig. 4.**
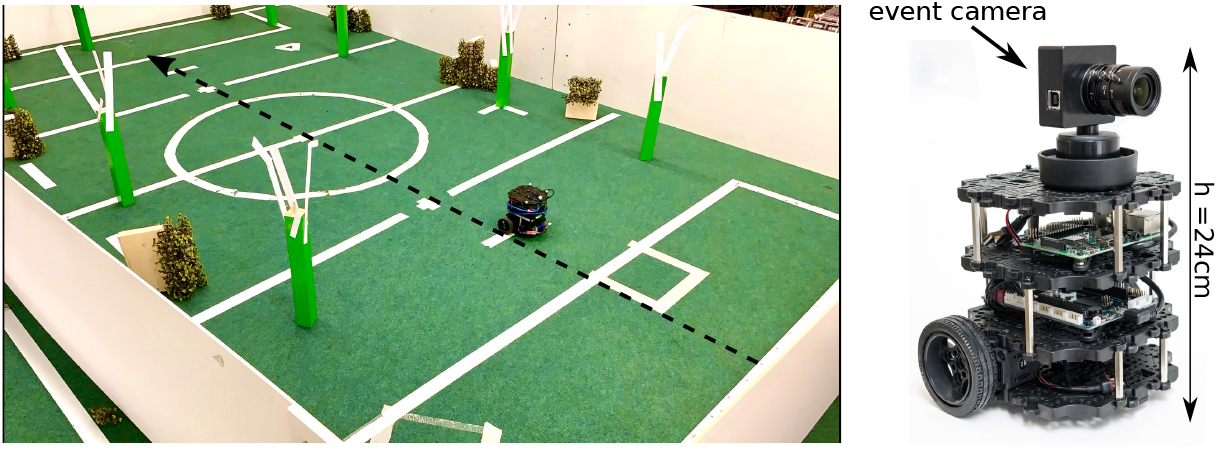
Test arena and robot hardware. The test arena (5 *m* × 3 *m*) consists of clean background and artificial vegetation randomly placed around robot pathways.

### Experiment 1: Learning route segments

We trained the network with segments of visual motion from a 4 m route (black trajectory in figure 4), then tested the trained network for a replay of the whole route. The training segment varies in length (0.4 m to 2 m) and in location (taken from the start or middle of the route). The MBON membrane potential (input from KCs) and spiking rate during each test are plotted in figure 5.

**Fig. 5.**
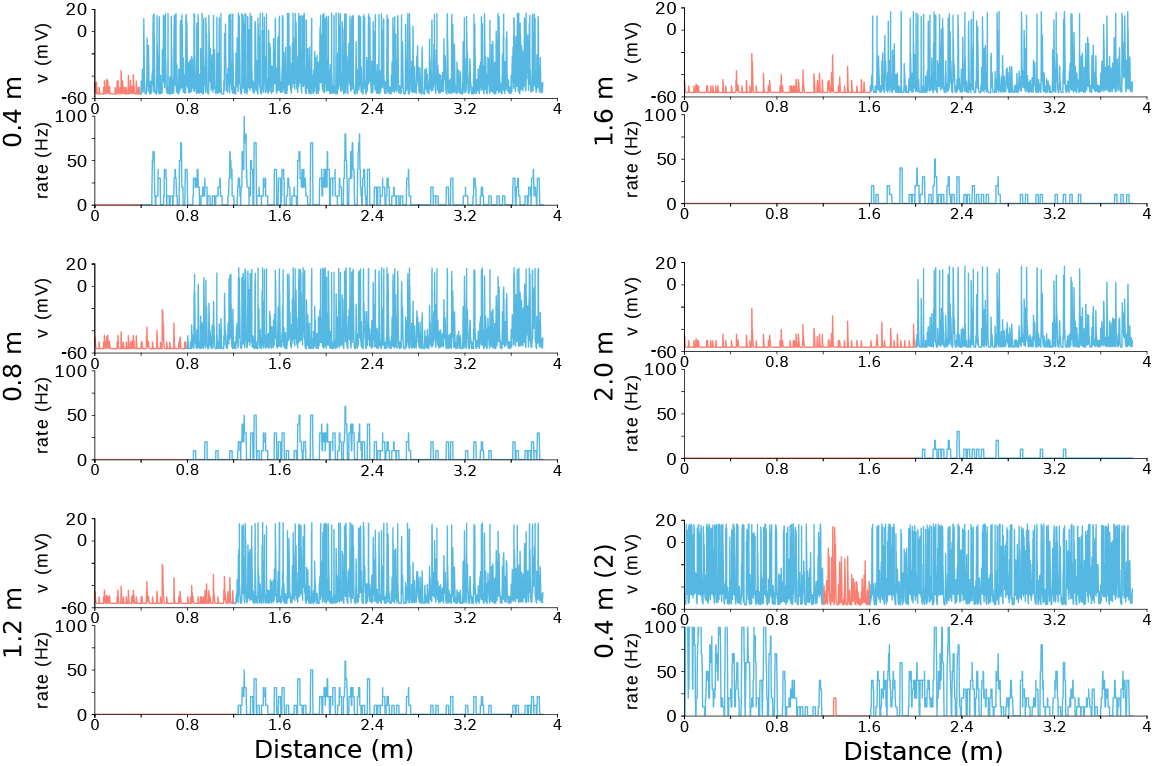
MBON membrane potential and spiking rate plotted against travelled distance. Pink: learned route segment. Blue: unlearned segment.

Note that KC activity, and hence MBON output, is partly influenced by the density of visual cues, so will vary even in unlearnt segments. However, it is clear that for each learned segment, the MBON spiking rate drops dramatically, indicating this part of the route is recognised as familiar by the network. After learning the visual flow over some travelled distance, the network memory capacity will be reached. This is because the longer the segment, the more KC-KC inhibition will be strengthened, and hence the MBON output will tend to reduce across all input patterns, trained or untrained. This can be seen in figure 5 ‘2.0 m’, where the low rate of MBON spiking overall could make it difficult to distinguish learned (0-2 m) from novel (2-4 m) visual input. We return to this issue of memory capacity in the discussion. Also note that the drop in MBON input is less dramatic when the learned segment was taken from the middle of the route (figure 5 ‘0.4 m (2)’). This reflects the effect of ‘internal noise’ in the network dynamics: in the test, the network state at the start of the learned segment (1.2 m) depends on the preceding input, and hence differs from the initial (resting) state used when training with that segment. This affects the relative timing of the subsequent KC spikes, despite the same input during the segment, so the KC pattern is not identical. Nevertheless the segment that was learned can be easily distinguished.

### Experiment 2: Adding noise

Experiment 1 showed how the ‘internal noise’ can affect the familiarity. Here we explicitly test the effect of adding external noise to the input, by replacing some camera events with randomly generated events (i.e. in different pixels) at the original event time. We introduce a novelty 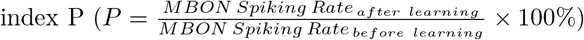 as a measure of the drop in MBON activity across a segment of the route. The results in figure 6 show that a noise level of 5% or more (equivalent to changing 1 input event every 1 ms) results in output for the learned segment that can no longer be distinguished from the unlearned segment, but for noise of 2% or less, the learned segment can be recognised.

**Fig. 6.**
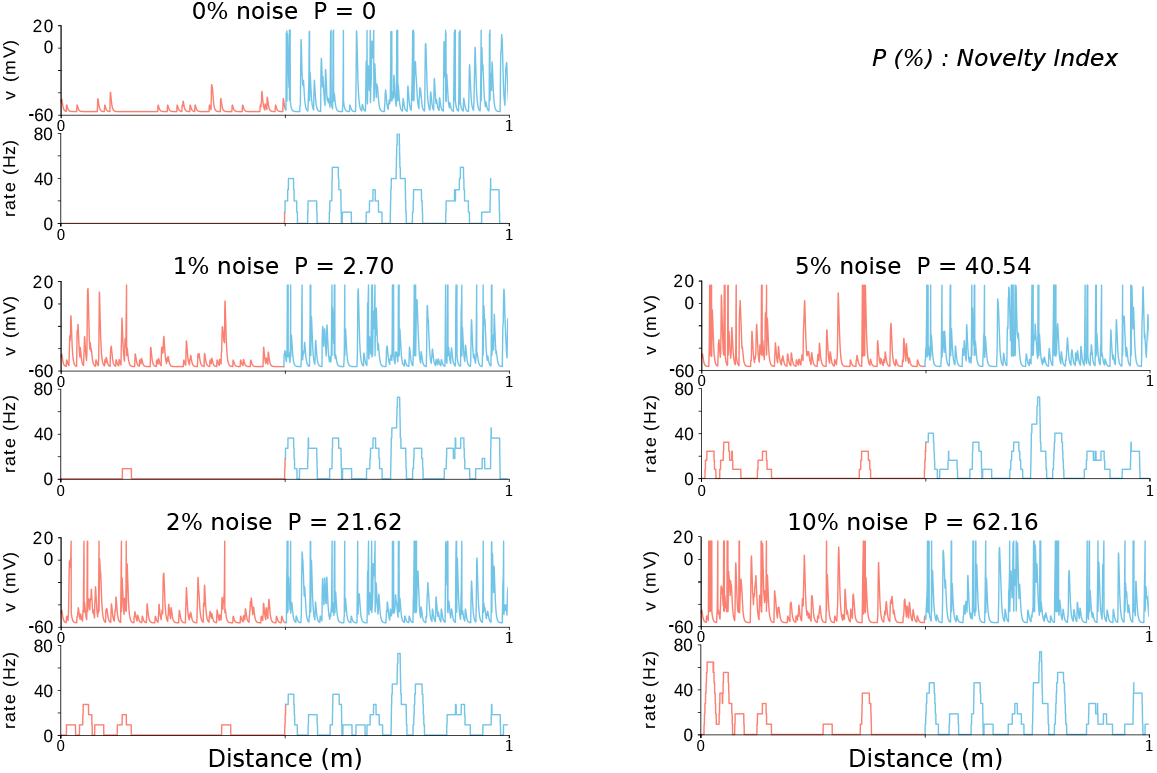
MBON membrane potential and spiking rate plots in experiment 2 noise test. Visual motion flow on learned routes (pink) is tested with adding 1% to 10% noise.

### Experiment 3: Off-route

A successful navigation model needs to not only recognise when it is on a familiar route, but also to be able to correct for deviations from the correct route. In previous image matching models, this is achieved by using gradient of familiarity around the positions where images are stored. Thus, ideally, the output of our network should remain low for small displacements (when the agent is almost but not quite on route) and increase as it is moved away from the route.

We firstly tested in a large arena (5 *m* × 3 *m*, figure 4) with relatively large displacements (10-50 cm or more parallel, or travelling 40 cm at angles 5-60 degrees, figure 7). Before learning, the MBON firing rate varies between 200 Hz to 400 Hz. The variation is due to the unevenly distributed vegetation (see figure 2 and figure 4). After learning, testing the network with displacements parallel to the route, or at increasing angles to the route shows a ‘valley’ of novelty, dropping to 0 on the learned route.

**Fig. 7.**
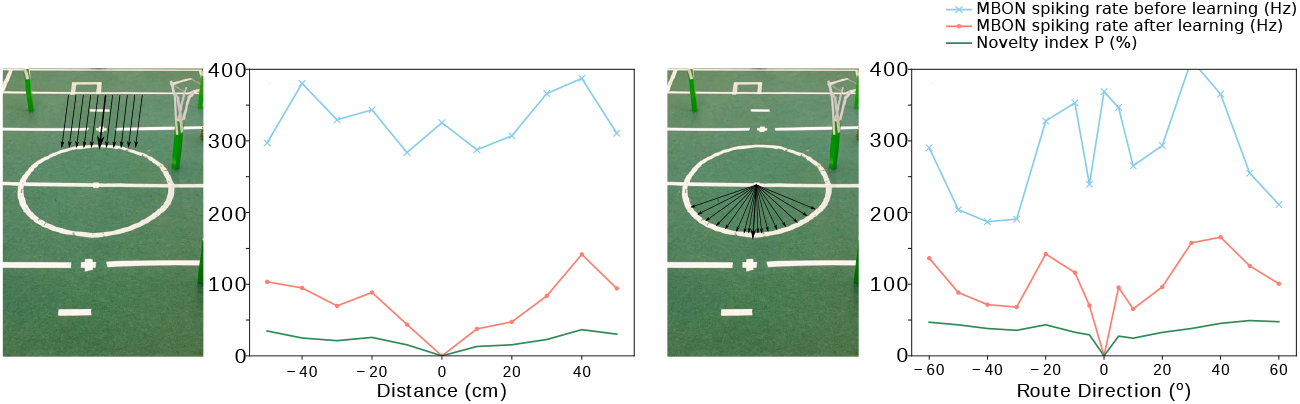
Off-route tests. Averaged MBON spiking rate before (blue) and after (pink) learning are plotted, together with novelty index P (green) for different displacements parallel or at angle to the learned route segment.

We then tested smaller displacements in a 1.5 *m* × 1 *m* arena (figure 8, a). The results in figure 8 show a similar trend, with a low novelty index for the learned route (0,0), increasing with 2-12 cm parallel displacement (W) or 2-6 cm backwards displacement (S). The angular change has a stronger effect in increasing novelty even from 10 degrees. Note that in most image-matching models, it is assumed that rotation on the spot will reveal the best matching direction. Here, it would not be practical for the robot to move along each angular segment to find which best matches the visual flow experienced on the route. We return to this issue in the discussion.

**Fig. 8.**
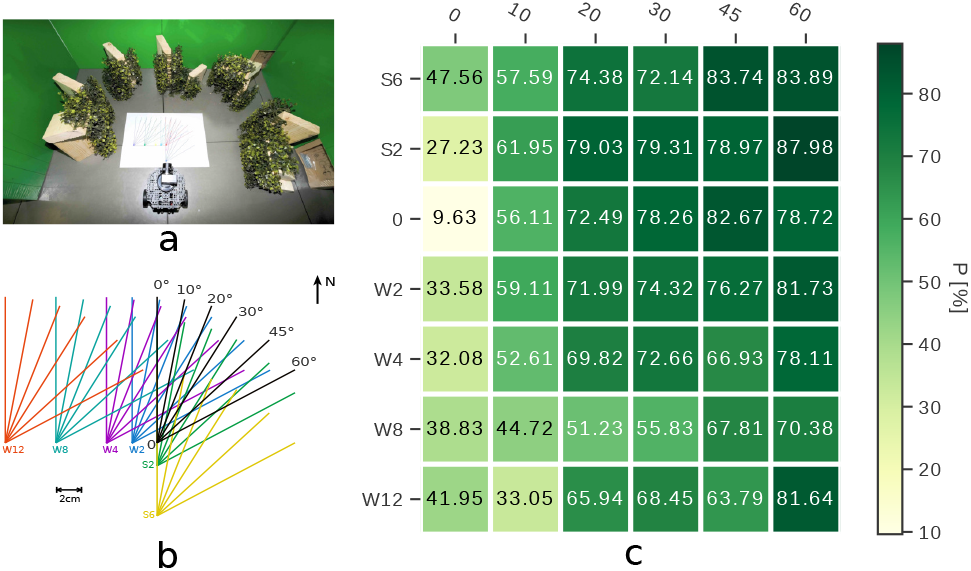
Off-route test in a smaller arena with denser vegetation. Map in (b) shows the spatial distribution of learned and tested routes in (a) (W: west, S: south). (c): Novelty index P of all testing routes

### Experiment 4: Sequence

Insects arriving a familiar location but in an unexpected order can trigger scanning and learning behaviour [7], suggesting sequence plays a role in visual memory. However, this could imply memory of ordered static images or of continuous motion flow; provided some temporal information about the input pattern is stored. Here we mimicked a recent experiment which tested sequence memory in ants by looking at the number of scans they made (indicative of unfamiliarity or confusion) when they experienced two familiar route segments in the correct order or in a novel order. After training the network on 1.5 m of a 2 m route (in figure4 arena), the learned motion flow is: 1) chopped into 3 parts and the order rearranged; 2) reversed in direction, i.e. what would be experienced by the robot going backwards from the end to the start of the route, without changing heading direction. The result (figure 9) is that MBON activity rises at the novel junctions formed between familiar segments and decreases again as the familiar segment is traversed. If MB activity controls scanning behaviour in ants, this would explain the observed ‘sequence’ effects in ant behaviour, i.e. that they scan when route segments occur out of order. Reversing the input sequence causes dramatic increase of novelty.

**Fig. 9.**
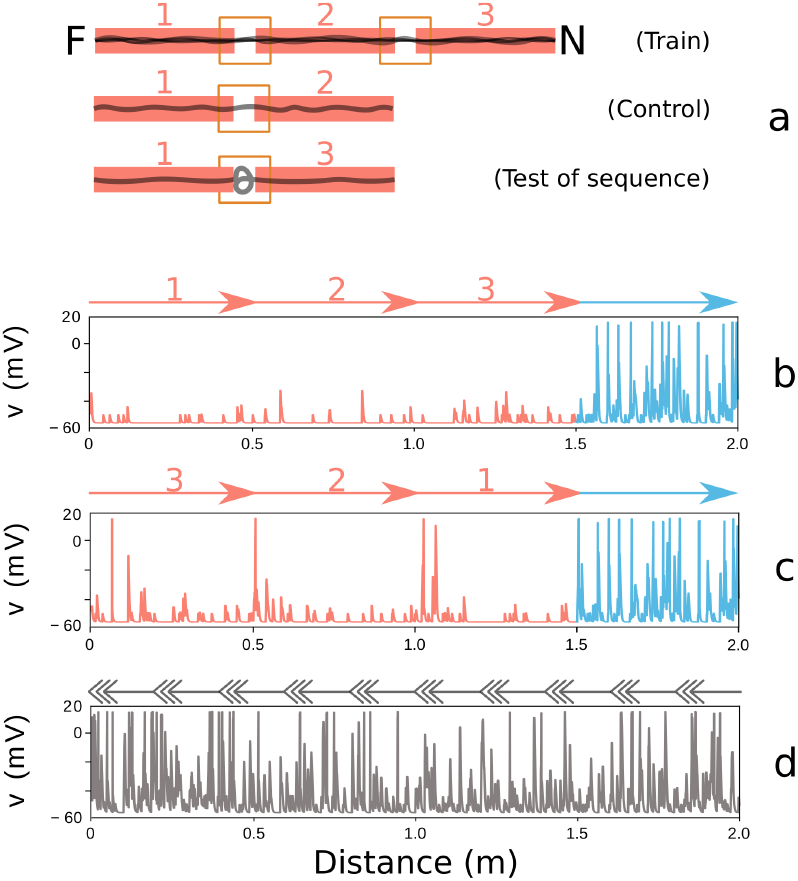
Sequence test in ants (based on [Schwarz et al, under review]) and results of experiment 4. a: Ants learned a route from F (feeder) to N (nest) which started with a distinctive channel (segment 1). Ants tested with the novel sequence [1,3] stopped and scanned at the unexpected junction between two familiar segments, whereas control ants [1,2] ran straight. b: the model produces a low output throught the correct sequence, c: the model spikes more at junctions when the sequence is reordered, d: a completely reversed input (running backward) is not recognised.

## 4 Discussion

We have presented a plausible neural mechanism by which route memory based on continuous visual motion can be stored. This is based on the insect MB connectome, and postulates a role for the recently observed, but functionally unexplained, KC-KC interconnections. In our spiking neural network model, the KC-KC interconnections can learn a spatio-temporal pattern using the STDP learning rule. Robot tests in two indoor environments have shown that this model can distinguish familiar from novel visual motion patterns, with some robustness to noise and small displacements. Route learning based on motion inherently encodes short sequences, and we show the model output is consistent with observed behaviour of ants when familiar segments of a route occur in a novel sequence. Our model has not yet been tested in more complex environments, but the next step will be to verify its robustness and generalisability in a natural environment.

The current model has a limited memory capacity (see experiment 1), but some simple modifications could increase the capacity: tuning the parameters to make KC firing sparser and thus slow the accumulation of inhibition; or simply increase the number of KCs. The size of MB in our model (only 10,000 KCs) is smaller than for insect navigators (e.g. the honeybee mushroom body has about 368,000 KCs [8]). Also, our model does lifetime learning of all incoming information, with no selection or forgetting. Learning in insect the MB is likely to be more sophisticated and efficient in dealing with redundancies in the sensory stream. This might also enhance robustness, as it seems likely that a repeated run on the same route in realistic conditions may easily exceed the 2% noise tolerance shown in Experiment 2. Introducing additional smoothing and motion processing layers between the event-based camera output and the MB input is also likely to be helpful. In the insect there are multiple neural layers of visual processing. Modelling these and other properties of the visual pathway is one of the obvious next steps for this work.

Another required step is to design a navigation strategy which can utilise the output of this model to generate proper route following patterns. Our model explains how an insect would know it is on route, but not how it would re-find the route. The novelty index results in Experiment 4 suggested that getting closer to the route should provide some drop in novelty, so an oscillation strategy as suggested in [10] could be effective, and will be explored in future work.

